# Regional epithelial architecture and spatial distribution of T and B lymphocytes in the human fallopian tube

**DOI:** 10.64898/2026.03.13.711514

**Authors:** Filippa Bertilsson, Feria Hikmet, Hjalte Sveidqvist, Michaela Einarsson, Theodora Kunovac Kallak, Matts Olovsson, Loren Méar, Cecilia Lindskog

## Abstract

The human fallopian tube plays a critical role in reproduction, yet its structural organization and immune landscape remain incompletely characterized. Here, we analyzed tissue from women of reproductive age across three anatomical regions (isthmus, ampulla, and fimbriae) throughout the menstrual cycle. By using antibody-based imaging together with automated image analysis, the epithelial thickness and spatial distribution of T and B lymphocytes was assessed. No significant differences in epithelial thickness were observed between proliferative and secretory phases. However, significant regional variation was identified, with the epithelium thickest in the isthmus and thinnest in the ampulla. Both CD8A^+^ T lymphocytes and CD20^+^ B lymphocytes were present throughout the fallopian tube, with their abundance strongly correlated across patients. Spatial analysis further revealed that both lymphocyte populations were preferentially localized to the mucosal compartment adjacent to the lumen, and intraepithelial B lymphocytes were consistently observed. Together, these findings provide new insight into epithelial organization and immune cell distribution in the human fallopian tube, highlighting the complexity of the tubal immune microenvironment and its potential relevance for reproductive biology.

## INTRODUCTION

The human fallopian tubes (FT) are paired muscular organs extending distally from the uterus toward each ovary, and serve as passage for the oocyte, sperm, and the early embryo. The FT exhibits distinct anatomical and morphological differences along its length, and can be divided into four regions: the uterotubal junction (also called the intramural segment), which connects the tube to the uterus; the isthmus; the ampulla; and the infundibulum, which contains the fimbriae, the fringed end of the tube that opens into the peritoneal cavity^1,2^. Each of the regions has distinct cellular compositions and physiological roles during fertilization. The isthmus, with an average length of 2-3 cm, has a narrow lumen surrounded by a thick muscular layer and regulates the passage of spermatozoa into the tube as well as the transport of the developing embryo into the uterus for implantation (Figure 1A)^1^. The ampulla, the longest part of the tube (5-8 cm), has a thinner muscular layer than the isthmus, and is the most common site of fertilization. Its luminal diameter can reach up to 1 cm (Figure 1A). The infundibulum, which has a trumpet-like shape, terminates in a fimbriated end responsible for oocyte capture and transport toward the ampulla for fertilization (Figure 1A)^1^.

**Figure 1.**
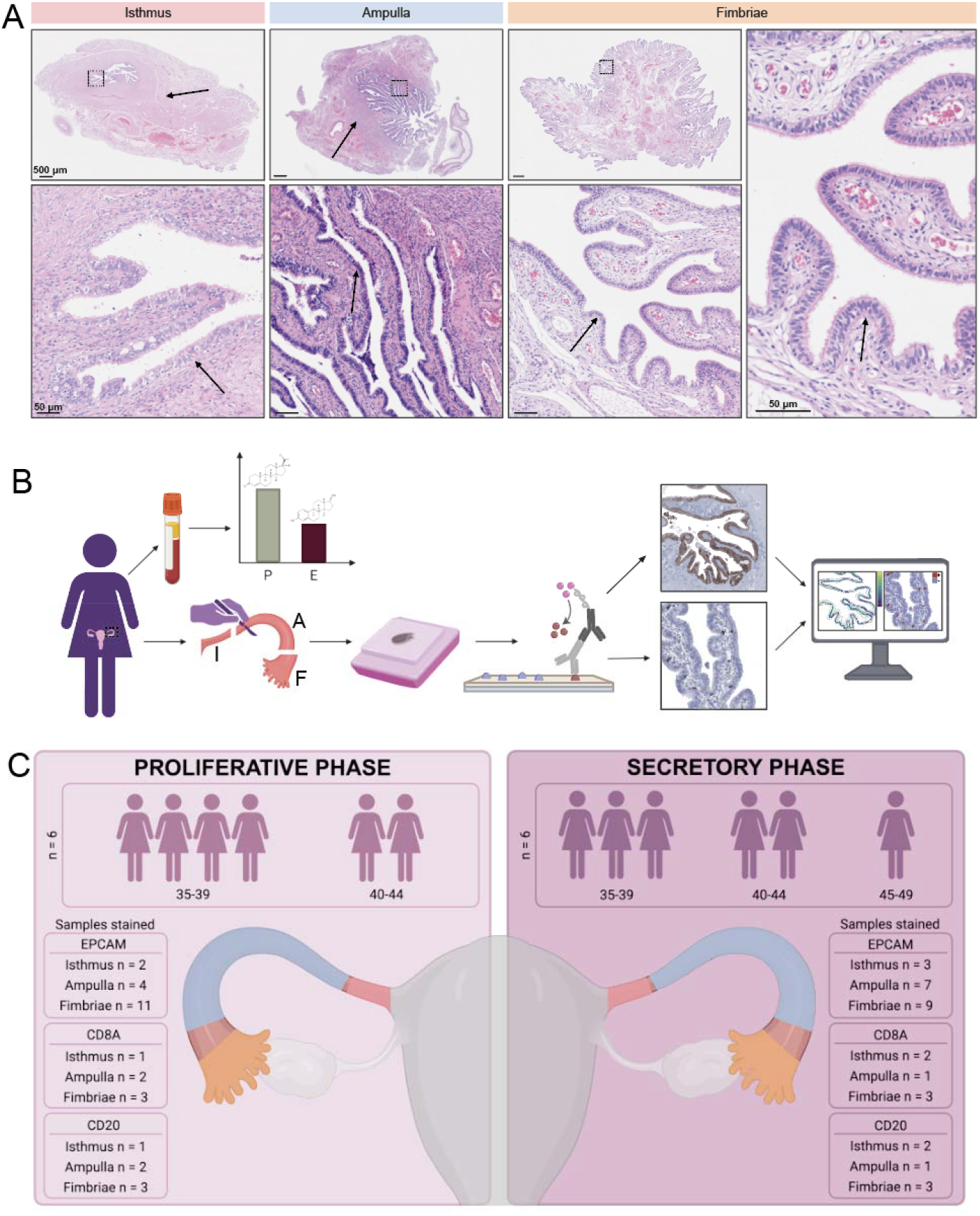
Fallopian tube histology and experimental design. **A) Tubal histology.** H&E staining of the isthmus, ampulla, and fimbriae. Arrows in the top row point at muscularis, while arrows in the bottom row and the right image point at the epithelium. **B) Workflow**. Fallopian tubes from women of reproductive age were collected along with blood samples to analyze progesterone (P) and estrogen (E) levels. The fallopian tubes were dissected into isthmus (I), ampulla (A), and fimbriae (F), embedded as FFPE blocks, and stained with immunohistochemistry. Images were analyzed to quantify epithelial thickness and immune cell abundance. **C) Cohort**. Six patients in the proliferative and six in the secretory phase were included in the study. Age ranged from 35-44 in the proliferative group and 35-46 in the secretory group. In total, 36 samples were stained with anti-EPCAM (17 proliferative and 19 secretory). For CD8A and CD20 staining, six samples per group were analyzed, with the same samples stained with both anti-CD8A and anti-CD20 antibodies.

The lumen of the FT is lined by an epithelial mucosa consisting of secretory and ciliated cells arranged in a columnar layer^3^. The height of this epithelium has been shown to fluctuate in response to circulating progesterone and estrogen levels^4^. Other cell types have also been reported within the epithelium, including Peg cells, which are thought to have stem cell properties^5^, and intra-epithelial T lymphocytes^6^. However, the exact identity and function of these T lymphocytes in the healthy FT remains unclear, and the broader composition and functional roles of immune cell populations in the human fallopian tube are still poorly understood^7^. Several studies have investigated immune cell populations in the FT, yet many report contradictory findings regarding the most abundant immune cell type and their fluctuations across the menstrual cycle. Although immune cells belonging to both the innate and adaptive responses have been identified ^8–10^, most studies have focused on T lymphocytes and macrophages. In contrast, relatively few studies have examined B lymphocytes, and some have failed to detect them entirely^7^.

Beyond the healthy FT, immune cells have been studied in pathological conditions such as ectopic pregnancy, where T lymphocytes have been localized at the implantation site within the tube using CD3^8^ or CD8^9–11^ markers. Immune cell populations have also been investigated in hydrosalpinx, where CD4+ T lymphocytes were reported to be more abundant than CD8+ lymphocytes^12^, indicating a shift compared with the immune profile observed in the healthy FT. B lymphocytes, macrophages, monocytes, and neutrophils have also been detected in hydrosalpinx ^13–15^, and in particular macrophages have been studied in relation to fertility within the FT^13^. The presence of macrophages, monocytes, and neutrophils in hydrosalpinx has been suggested to contribute to reduced fertility in affected patients^12^. The potential impact of immune cells in the FT in fertility remains however unclear. A better understanding of the cell populations of both the innate and adaptive immune systems in the FT is therefore essential to elucidate their physiological roles in reproductive function^7^.

The aim of this study was to characterize the human FT epithelium and immune repertoire in women of reproductive age across different anatomical regions and throughout the menstrual cycle. Using immunohistochemistry (IHC) and quantitative image analysis (Figure 1B), we were able to provide new insights into epithelial thickness and the cellular complexity of the FT. Together, these observations highlight the need for a more comprehensive understanding of the fallopian tube immune landscape and its interaction with the epithelium. Here we provide further insights in the spatial relationship between epithelial and immune cells to provide a critical foundation for future studies of FT function in health and disease.

## METHODS

### Sample collection and preparation

All samples were collected and anonymized at Uppsala University Hospital (Uppsala, Sweden) in accordance with ethical approval from the Swedish Ethical Review Authority (2008-046). Twelve women of reproductive age with regular menstrual cycle were included, undergoing surgery for either uterine myoma or sterilization. Blood samples were collected prior to surgery to measure progesterone and estrogen levels and confirm the menstrual phase (proliferative phase n=6, and secretory phase n=6). Hormone serum levels were measured at the SWEDAC accredited Clinical Chemistry Department at Uppsala University Hospital. Five women in each group had previously been pregnant at least once. Collected tissue samples were divided into the distinct anatomical regions (isthmus, ampulla, and fimbriae). A total of 36 samples (Appendix Table 1) were assembled into individual formalin-fixed and paraffin-embedded (FFPE) tissue blocks. FFPE blocks were sectioned into 4 *µ*m sections using a waterfall microtome (Microm H355S, ThermoFisher Scientific, Fremont, CA). Sections were mounted onto SuperFrost Plus slides (Thermo Fisher Scientific, Fremont, CA) and air-dried at room temperature (RT) overnight before being baked at 50°C for approximately 16 hours. Slides were then stored at -20°C until use.

### Immunohistochemistry (IHC)

Before immunohistochemical (IHC) staining, slides underwent pre-treatment including de-paraffinization, rehydration, and heat-induced antigen retrieval (HIER) as previously described^14^. IHC staining was performed at RT in darkness using a LabVision Autostainer 480S (Thermo Fisher Scientific, Freemont, CA). Primary antibodies were used to label epithelial cells (anti-EPCAM, M0804, Agilent, Santa Clara, United States 1:50), CD8+ T lymphocytes (anti-CD8A, HPA037756, Atlas Antibodies, Stockholm, Sweden, 1:800), and CD20+ B lymphocytes (anti-CD20, HPA014341, Atlas Antibodies, Stockholm, Sweden, 1:400). All three antibodies have been validated as part of the Human Protein Atlas workflow, with data and images available at www.proteinatlas.org. The validation includes IHC staining across 44 different normal tissues, thereby verifying the expected protein expression pattern with EPCAM expressed in specific epithelial cells, and CD8A or CD20 positive cells in lymph node mainly localized to paracortex or germinal centra, respectively. A quality control staining was performed to further validate the antibodies in thyroid, appendix, and an additional fallopian tube sample not included in the cohort (Appendix Figure 1). The staining protocol consisted of the following steps: Peroxidase Block (925B-04, Cell Marque, Rocklin, California, United States) for 10 minutes, incubation with primary antibody (anti-EPCAM, anti-CD8A, or anti-CD20) diluted in BrightDiluent (UD09-125, ImmunoLogic, MS Arnhem, The Netherlands) for 30 minutes, UltraVision Primary Antibody Enhancer (954D-32, Cell Marque, Rocklin, California, United States) for 10 minutes, HiDef Detection HRP polymer detector (Cell Marque, 954D-32) for 10 minutes, and finally DAB Quanto Chromogen (957D-31, Cell Marque, Rocklin, United States) diluted in DAB Buffer (957D-32, Cell Marque, Rocklin, California, United States) for 10 minutes. Slides were subsequently counterstained and mounted according to a previously described protocol^14^. Whole-slide images were acquired in brightfield at 40x magnification using an Akoya PhenoImager HT digital slide scanner (Akoya Biosciences).

### Image analysis

Images were analyzed using the open-source software QuPath (version 0.6.0)^15^. For epithelial thickness measurement, one region of interest (ROI) was manually annotated for each image based on the highest-quality epithelial layer. A classifier was trained to distinguish epithelium from the surrounding tissue and applied within each ROI to generate an object outlining the epithelial layer. A manual QC was done for each object to remove potential erroneous detections. Each object was exported as GeoJSON-files containing vector polygons and subsequently analyzed in Python (version 3.11.14) using Spyder (version 6). The Python analysis pipeline was based on a previously described method^16^, where per-point epithelial thickness measurements were extracted using euclidean distance transform.

For CD8A+ and CD20+ lymphocyte quantification, two anatomical layers were annotated where possible: the mucosa (including the epithelium and lamina propria), and the muscularis (including the surrounding muscular layers and peritoneum). Positive Cell Detection in QuPath was applied within each annotation to determine the total number of cells and the number of marker-positive cells. Results were exported as tables for downstream analysis. Density maps were performed using the “Density maps” function in QuPath.

### Data analysis

Statistical analyses and visualization statistics were performed in RStudio (version 4.5.2). Plots were generated using the package ggplot2 (version 4.0.1)^17^. Epithelial thickness was compared between menstrual phases and within each anatomical region (isthmus, ampulla, and fimbriae) using Welch two-sample t-test, applied with the rstatix package (version 0.7.3)^18^. To compare epithelium thickness across anatomical regions while disregarding the menstrual phase, a linear model was generated using the lmerTest package (version 3.2-0)^19^. An ANOVA was applied to the model to test the null hypothesis (H0) that mean epithelial thickness is equal across the isthmus, ampulla, and fimbriae, against the alternative hypothesis (H1) that at least one region differs (p < 0.05). Estimated marginal means (or adjusted means) were then calculated using the emmeans package (version 2.0.1)^20^ to identify pairwise differences between FT anatomical regions.

The abundance of CD8A+ and CD20+ lymphocytes across the anatomical regions and menstrual phases were due to the small sample size compared by applying a Wilcoxon rank-sum test using the rstatix package (version 0.7.3). P-values were adjusted with the Holm correction method. The association between CD8A+ and CD20+ cell counts within patients was evaluated using Spearman correlation with a significance threshold of p < 0.05. To assess whether the immune cells were more preferentially located near the luminal compartment compared with the surrounding muscular layers, a paired Wilcoxon signed-rank test was applied, with a significance threshold of p < 0.05.

### Data and code availability

All scripts will be available on GitHub (https://github.com/LindskogLab) at the time of publication. All images are found on figshare.com with identifier: https://doi.org/10.6084/m9.figshare.31860724

## RESULTS

### Epithelial thickness varies more across anatomical regions than across the menstrual cycle

FT samples from 12 women of reproductive age (37-46 years old) with regular menstrual cycles (Figure 1C) were used to study epithelial thickness and immune cell populations across anatomical regions and menstrual cycle phases. Samples from six women corresponded to the proliferative phase of the menstrual cycle and six to the secretory phase, as confirmed by serum hormone levels (Appendix Figure 2).

**Figure 2.**
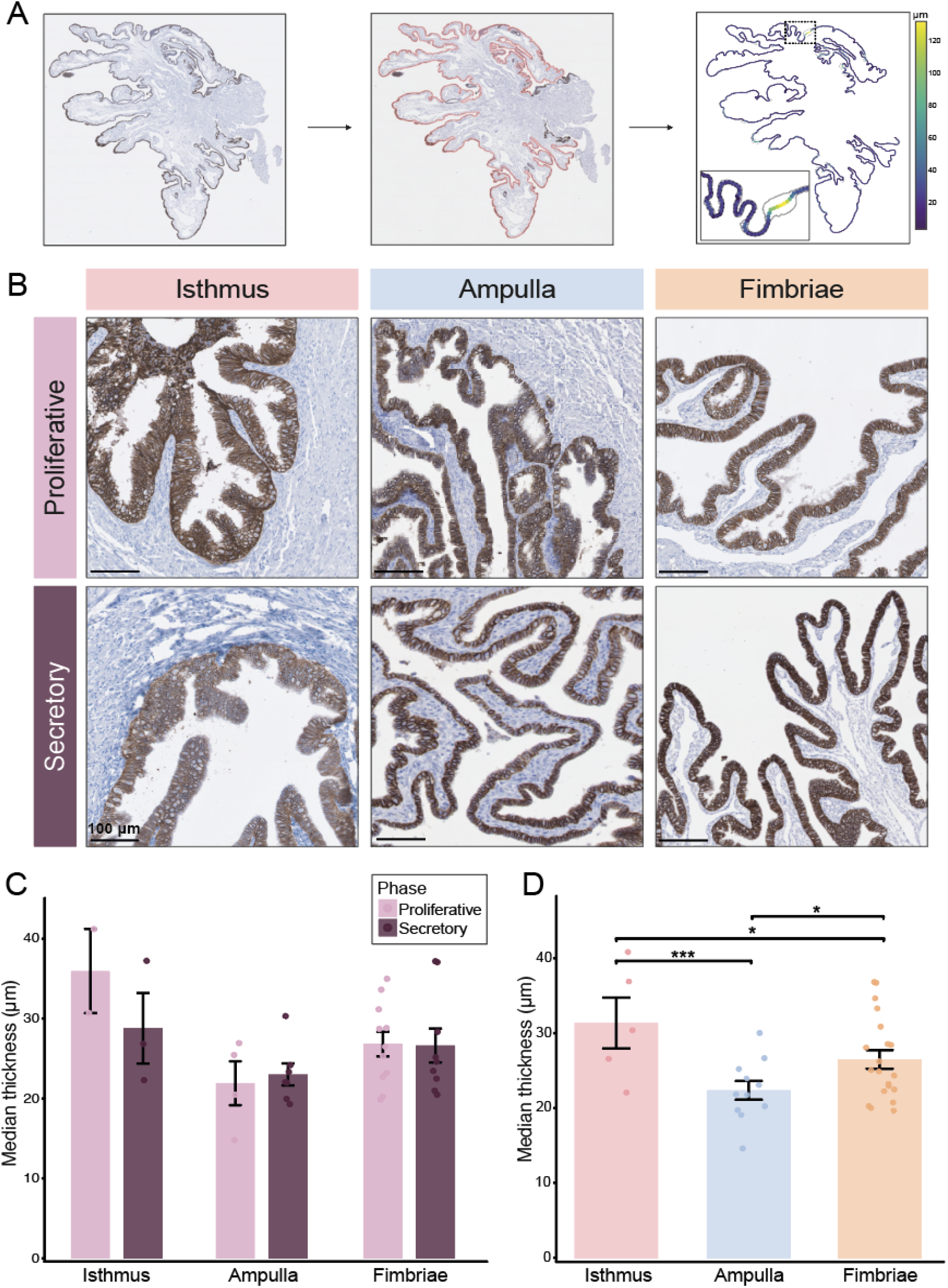
Epithelial thickness across the menstrual cycle and tubal parts. **A) Epithelial thickness measurement workflow**. Example illustrating how epithelial thickness was calculated. The epithelium was detected using image analysis, highlighted in pink (middle). Thickness measurements were then calculated across the epithelial layer at regular intervals, generating a thickness map in which thicker areas are shown in yellow and thinner areas in blue (right). **B) Immunohistochemical staining**. Representative images of isthmus, ampulla, and fimbriae, stained with anti-EPCAM antibody in both proliferative and secretory phase. Scale-bar = 100 µm in each image. **C) Epithelial thickness across the menstrual phases**. Calculated epithelial thickness of isthmus, ampulla, and fimbriae, in proliferative and secretory phase. Dots represent the median thickness of each analyzed image. No significant differences were observed between phases within any of the regions (isthmus: p = 0.393, ampulla p = 0.735, fimbriae p = 0.944). **D) Regional differences in epithelial thickness**. Epithelial thickness for isthmus, ampulla, and fimbriae with both menstrual phases combined. Dots represent the median thickness of each analyzed image. Significant differences were observed between the isthmus and ampulla (p = 0.0006) and between the isthmus and fimbriae (p = 0.0176). The fimbriae were also significantly thicker than the ampulla (p = 0.0218).

To compare epithelial thickness across tubal regions and menstrual cycle phases, samples from all patients (Figure 1C) were stained with IHC using an anti-EPCAM antibody (Figure 2B). EPCAM is expressed across all epithelial cells, enabling a clear visualization of the epithelial thickness. For each image, the epithelium was detected and outlined using automated image analysis, generating segmented epithelial objects with thickness measurements calculated across the epithelium (Figure 2A). Due to substantial intra-sample variability caused by the complex 3D structure of the folded mucosa, the median epithelial thickness was used for each sample (Appendix Figure 3). First, we examined whether epithelial thickness differed between proliferative and secretory phases within each tubal region. No significant differences were observed for any of the regions (isthmus: p = 0.393, ampulla: p = 0.735 and fimbriae: p = 0.944) (Figure 2C), suggesting that epithelial thickness does not significantly change across the menstrual cycle. When the menstrual phase was disregarded and all samples were analyzed together, significant regional differences however emerged. The epithelium was significantly thicker in the isthmus than in the ampulla (p = 0.0006) or the fimbriae (p = 0.0176) (Figure 2D). Comparing ampulla and fimbriae showed that the epithelium in fimbriae was thicker (p = 0.0218), thus concluding that the epithelium in the ampulla is thinnest within the FT (Figure 2D). Together, these results indicate regional differences with the epithelial thickness being greatest in the isthmus and lowest in the ampulla. Notably, epithelial thickness also showed a larger variability in the isthmus compared with the ampulla and fimbriae.

**Figure 3.**
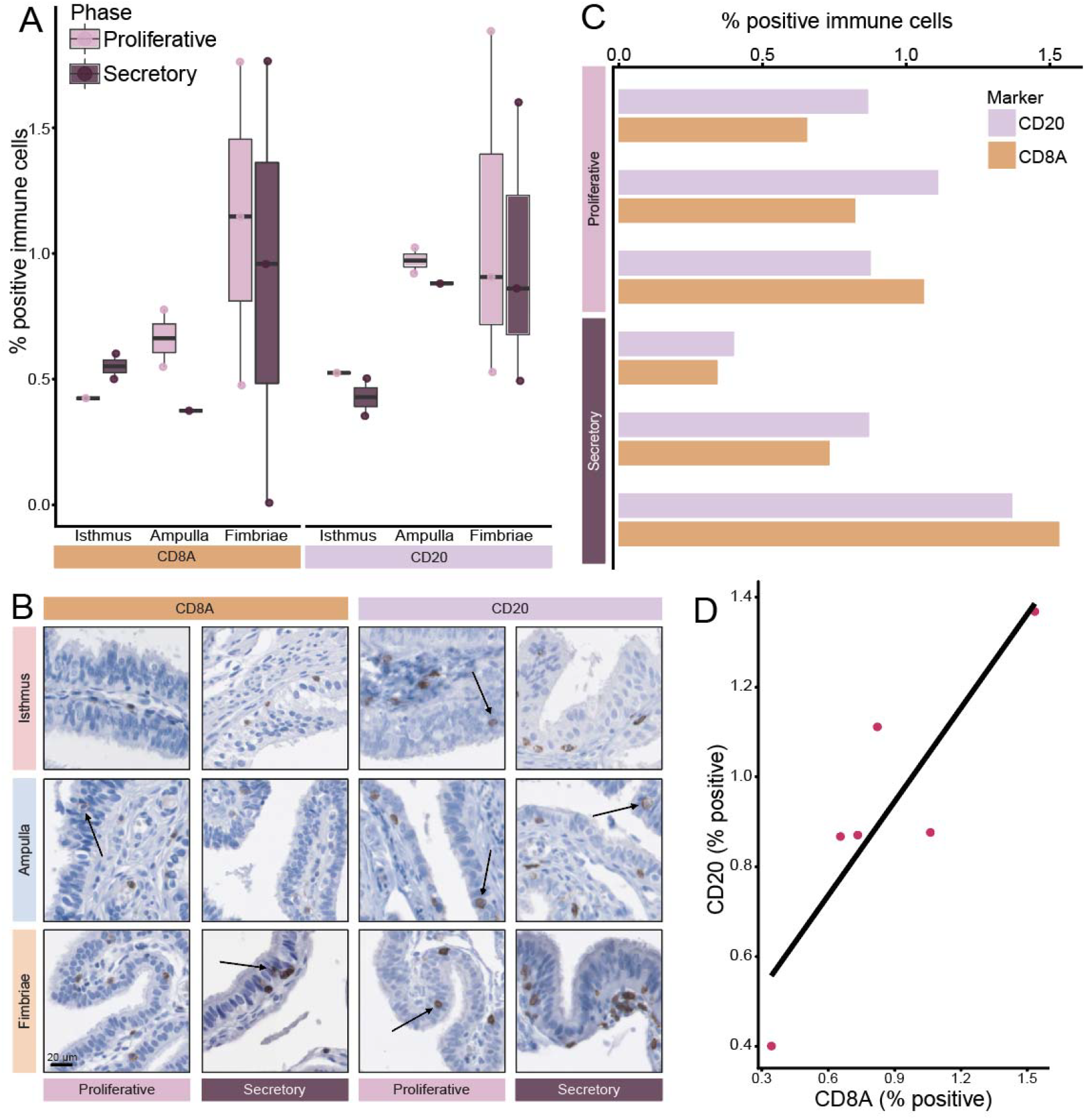
Distribution of T and B lymphocytes across fallopian tube regions, menstrual phase and patients. **A) Immune cell abundance across regions and phases**. Percentage of CD8A+ and CD20+ cells in isthmus, ampulla, and fimbriae during proliferative and secretory phases. No significant differences were observed between phases within any region (CD8A+: isthmus p = 0.667, ampulla p = 0.667, fimbriae p = 1.0; CD20+: isthmus p = 0.667, ampulla p = 0.667, and fimbriae p = 0.7). **B) Immunohistochemical staining**. Representative images showing CD8A+ and CD20+ cells in each anatomical region across both menstrual phases. Arrows indicate lymphocytes located within the epithelium. **C) Immune cell abundance per patient**. Percentage of CD8A+ and CD20+ lymphocytes for each patient in the proliferative and secretory phases (two bars per patient). **D) Correlation between T and B lymphocytes**. Spearman correlation of matched CD8A+ and CD20+ lymphocyte counts across patients (r= 0.94, p = 0.017).

### Distribution and abundance of T and B lymphocytes in the human fallopian tube

Next, we examined whether T and B lymphocytes, identified by anti-CD8A and anti-CD20 IHC staining respectively, varied across menstrual phase and tubal regions. Notably, all analyzed samples (Figure 1C) contained both CD8A+ and CD20+ immune cells. The largest variation in immune cell abundance was observed in the fimbrial region for both cell types however, no significant differences were detected between anatomical region or menstrual phases (Figure 3A). Since regional or phase-dependent differences were observed, immune cell abundance was further analyzed at the patient level. Interestingly, we detected high interpatient variability, with proportions ranging from <0.5% positive cells in some patients to >1.5% in others (Figure 3C). Despite this variability, a strong correlation between CD8A+ and CD20+ cell abundance was observed within patients (r = 0.94, p = 0.017), indicating coordinated T and B lymphocyte levels in the FT (Figure 3D).

### Spatial distribution of T and B lymphocytes within the human fallopian tube

Spatial analysis revealed that both CD8+ and CD20+ lymphocytes were frequently located beneath the epithelium across both the different tubal regions and the menstrual phases (Figure 3B). We were able to observe migration of both CD8A+ and CD20+ lymphocytes toward the lumen (Figure 3B). To determine whether immune cells preferentially localized near the epithelial compartment, immune cell density was compared between the mucosal layer and the surrounding muscularis. A significantly higher density of CD8A+ cells was observed near the lumen compared with the muscular layers (p = 0.042) (Figure 4A). A similar pattern was observed for CD20+ cells, with an even stronger enrichment in the mucosa (p = 0.00049) (Figure 4B). These results indicate that both T and B lymphocytes preferentially localize in proximity to the mucosal compartment. Density maps visualization further confirmed these findings, highlighting areas with increased immune cell density in the mucosa compared to surrounding muscularis (Figure 4C). Although lymphocytes were present in both annotated regions, their density was consistently higher in mucosal regions adjacent to the lumen.

**Figure 4.**
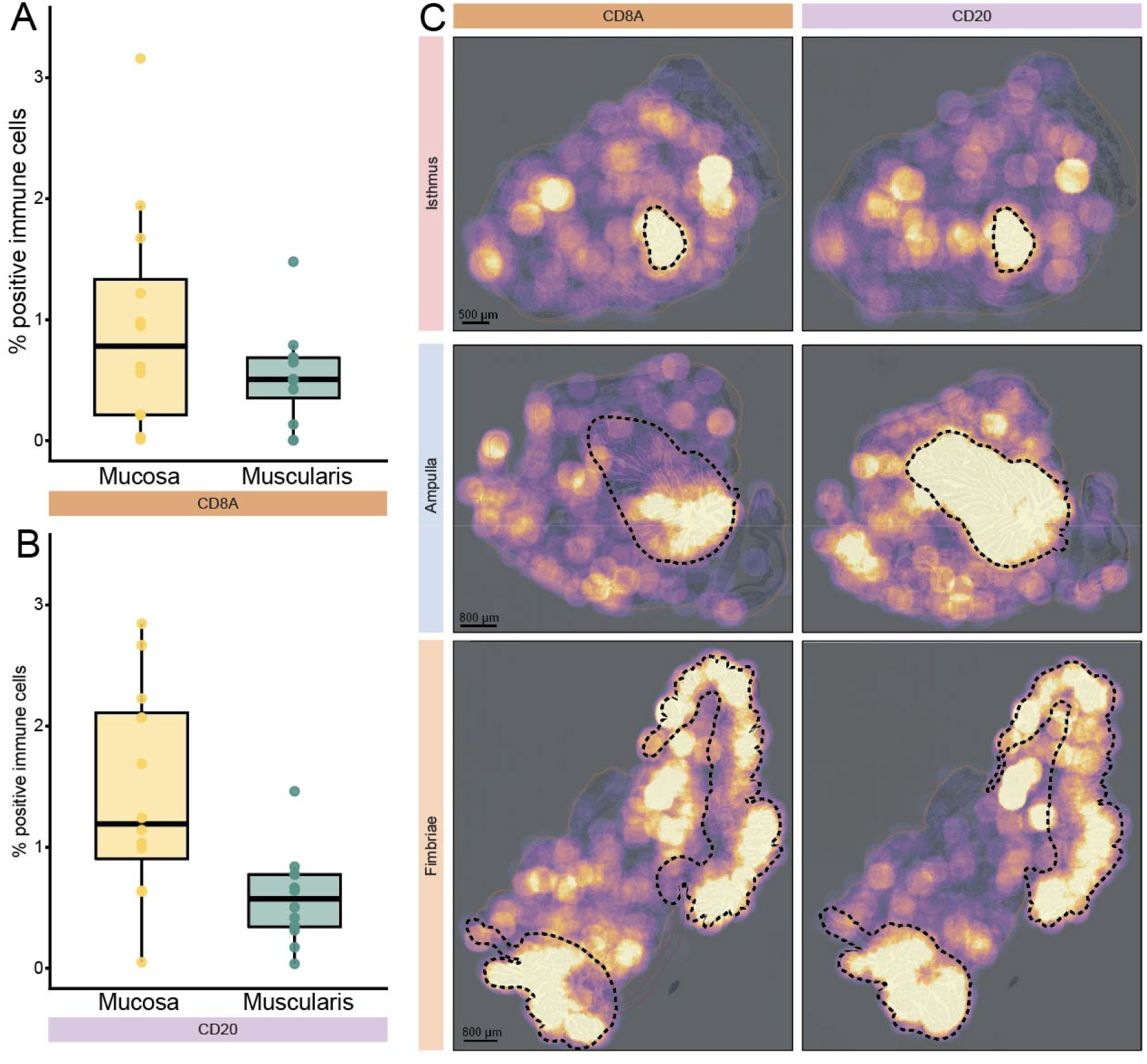
Spatial distribution of T and B lymphocytes in the fallopian tube. **A) Spatial location of CD8A+ lymphocytes.** Comparison of the percentage of CD8A+ cells located in the mucosal compartment (epithelium and lamina propria) versus the surrounding muscular layers. Statistical significance was assessed using a paired Wilcoxon signed-rank test (p = 0.042). **B) Spatial location of CD20+ lymphocytes**. Comparison of the percentage of CD20+ cells located in the mucosal compartment versus the surrounding muscularis. Statistical significance was assessed using a paired Wilcoxon signed-rank test (p = 0.00049). **C) Density maps**. Representative density maps showing the spatial distribution of CD8A+ and CD20+ cells in isthmus, ampulla, and fimbriae across proliferative and secretory phases. Dotted lines indicate annotated mucosal areas outlining the luminal compartment, including the lamina propria.

## DISCUSSION

Considerable effort has been devoted to studying the fallopian tube (FT) epithelium and immune cell populations associated with fertility in both health and disease. The present study provides additional insight into the structural organization and immune landscape of the human FT. First, we examined whether epithelial thickness fluctuated across the menstrual cycle by comparing proliferative and secretory phases within each anatomical region (isthmus, ampulla, and fimbriae). No significant differences between menstrual phases were detected. Cyclic changes in the FT have previously been reported, including alterations in epithelial thickness, epithelial protrusion, and ciliary beat frequency^1,4^. Fluctuations in progesterone and estrogen levels across the menstrual cycle are thought to influence protein production within the epithelium^3^, where secretory cells have been shown to release their contents at the time of ovulation, potentially increasing ciliary movement^21^. Although epithelial height has previously been reported to vary across the menstrual cycle^1^, we did not detect such changes in the present study. This discrepancy may reflect the limited sample size, and larger studies will be required to clarify whether subtle cycle-dependent differences exist.

In contrast, we identified significant regional differences in epithelial thickness. The ampulla exhibited a significantly thinner epithelium than both the isthmus and fimbriae, whereas the isthmus showed the greatest thickness. Although, regional variation in epithelia cell composition has been described^21^, this is, to our knowledge, the first demonstration of significant differences in epithelial thickness across FT regions. The thinner epithelium in the ampulla may relate to its relatively wide lumen (up to 1 cm in diameter)^1^ and functional role as the primary site of fertilization^22^, potentially facilitating gamete interactions. The ampulla has also been shown to secrete oviduct-specific glycoproteins in response to estrogen in sheep and cattle^23^, although whether similar mechanisms occur in humans and how they influence the tubal epithelium remain unclear. Further studies, incorporating transcriptomic and proteomic approaches will be required to fully characterize regional specialization of the tubal epithelium.

It is well established that the FT contains immune cells from both the innate and adaptive immune systems^24^. Several studies have reported T lymphocytes as the most abundant immune cell type in the FT ^24,25^, with CD8+ T lymphocytes representing the most common subset at both transcript and protein level ^24–26^. B lymphocytes, in contrast, have been reported either at lower abundance than T lymphocytes ^24^, or not detected at all ^24,25^. In the present study, however, we observed no significant difference in abundance between the T (CD8A+) and B (CD20+) lymphocytes across menstrual phases or tubal regions. Instead, a strong correlation between T and B lymphocytes abundance was observed across patients, suggesting coordinated levels of these cell populations within the FT. Interestingly, substantial variability in immune cell count was observed between patients, indicating a highly individualized immune cell landscape in the FT. How this variability relates to systemic immune status or reproductive outcomes remains to be determined, and further studies linking immune cell counts in the fallopian tube to other clinical parameters are clearly warranted.

Importantly, this study provides evidence of intra-epithelial B lymphocytes in the human FT, a finding not previously reported since previous reports have only described intra-epithelial T lymphocytes ^12,24,25^. While previous studies have localized B lymphocytes primarily to the lamina propria ^24,25^, our data indicate that they can also populate the epithelial compartment across both menstrual phases and regions. Given that B lymphocytes remain relatively understudied in the FT, these findings highlight a previously unrecognized component of the tubal immune environment and warrant further investigation into their functional roles.

Based on previous reports indicating that T lymphocytes are primarily located within the epithelium^25^, we investigated whether a similar spatial distribution could be observed across the isthmus, ampulla, and fimbriae. Quantification of immune cells in the mucosal and muscular compartment revealed a significantly higher abundance of T lymphocytes in the mucosa surrounding the lumen compared to the muscularis. A similar pattern was observed for B lymphocytes, which also showed a strong enrichment within the mucosal compartment. Density map analysis further supported these findings, demonstrating that both T and B lymphocytes were predominantly concentrated within the mucosa. Although small clusters of lymphocytes were occasionally observed within the muscular layers, most cells were located within the folded mucosa across all examined tubal regions. While lower immune cell counts in the muscular layers are expected, no previous study has performed a comprehensive quantification of the immune cell abundance across different histological layers.

This study has several limitations. First, immune cell identification relied on a limited set of markers, which may not fully capture the cellular heterogeneity ^7^. For example, rare CD20-expressing T lymphocytes have been described^26^ particularly in pathological context^27^, and potential marker overlap cannot be excluded. Nevertheless, antibody validation demonstrated staining patterns consistent with those expected in defined regions of lymphoid tissues. Second, the relatively small sample size limits statistical power. Future studies including larger cohorts, expanded marker panels, and multi-omics approaches will be essential to validate and extend these findings.

In summary, we investigated epithelial structure together with T and B lymphocyte population in the fallopian tube of healthy women of reproductive age. Although no changes in epithelial thickness were observed across the menstrual cycle, significant regional differences were identified between the isthmus, ampulla, and fimbriae, with the ampulla displaying the thinnest epithelium. In addition, we observed the presence of both T and B lymphocytes throughout the fallopian tube, with considerable inter-patient variability. Importantly, we provide evidence of intraepithelial B lymphocytes for the first time, highlighting the complexity of the tubal immune environment. This emphasizes the need for further in-depth research into different immune cell populations in the fallopian tube, and the potential role of the immune repertoire in normal reproduction of different tubal pathologies.

## ACKNOWLEDGEMENTS

We thank the patients who donated tissues for the research. We also thank the members of the HPA Tissue profiling team for supporting us in the project with antibody testing, IHC staining and imaging. The project was funded by the Swedish Research Council (2022-02742) and Cancerfonden (23 3003 Pj/25 4217 IA SIA).

