## Appendix Figures for "Regional epithelial architecture and spatial distribution of T and B lymphocytes in the human fallopian tube"

**Appendix Figure 1**

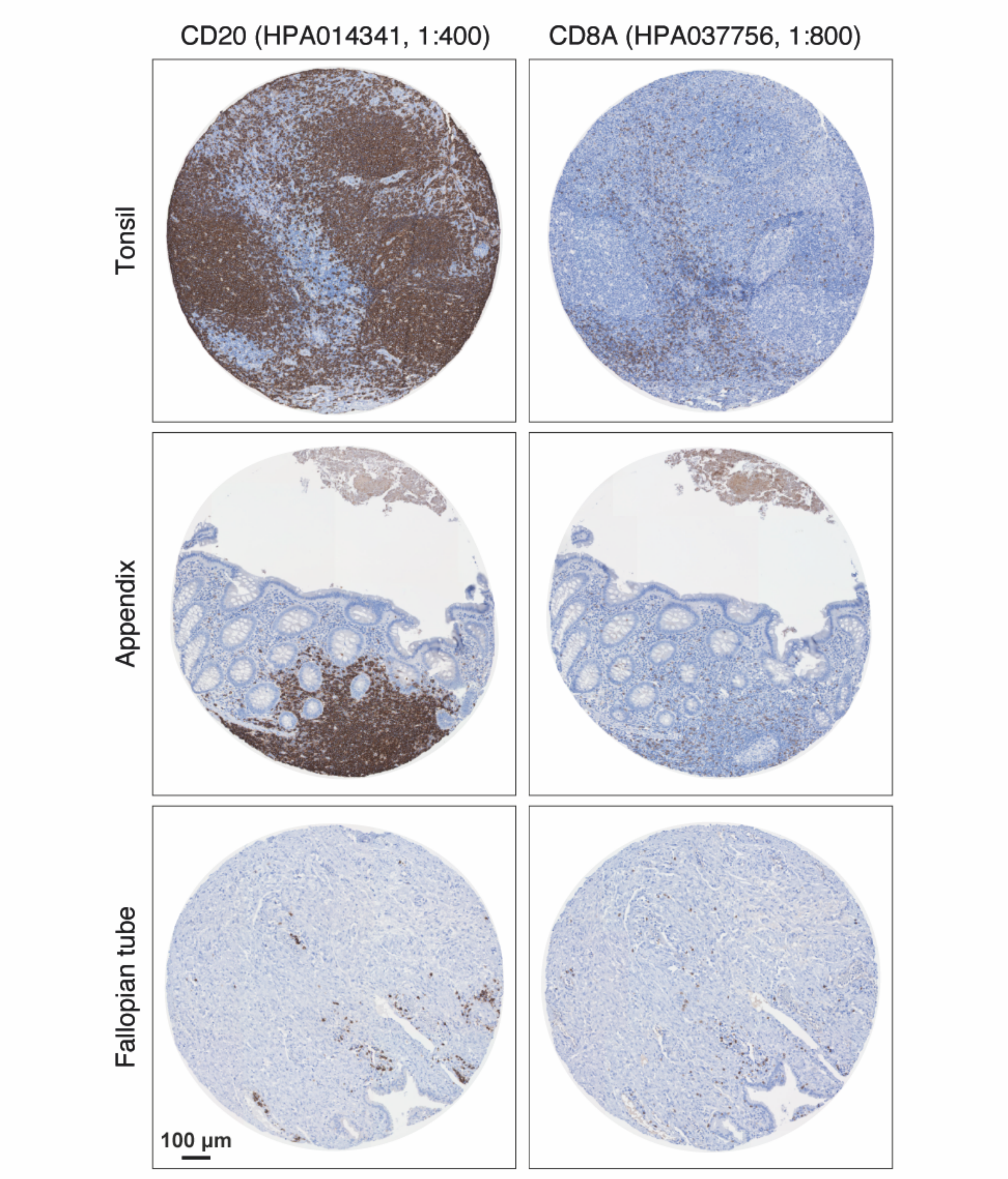


**Quality control (QC) of antibodies targeting B and T lymphocytes.** QC of the antibody targeting CD20 (left) and CD8A (right) in tonsil, appendix, and fallopian tube. The fallopian tube sample is not from the studied cohort.

**ppendix Figure 2**

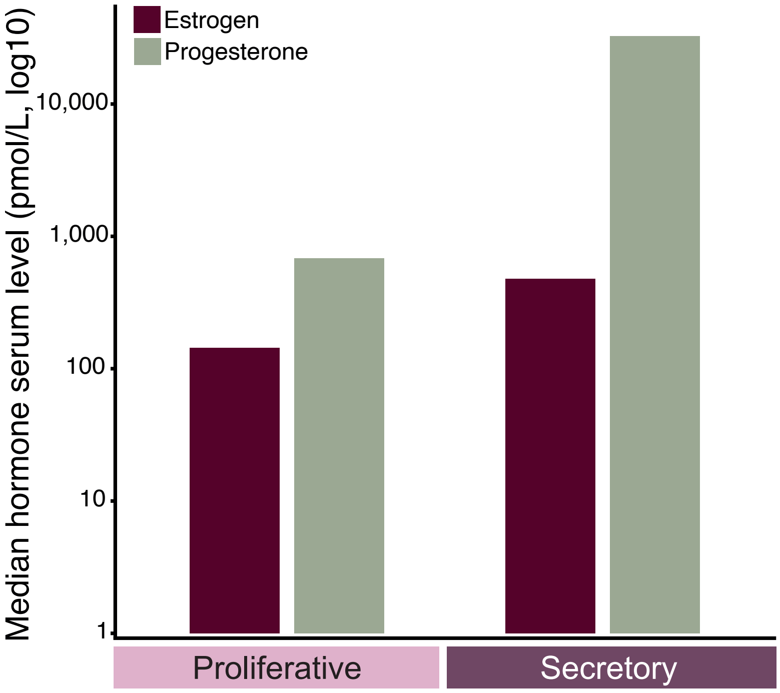


**Median hormone serum level (pmol/L).** The median estrogen and progesterone levels of the patients measured in serum on log10 scale**.**

**Appendix Figure 3**

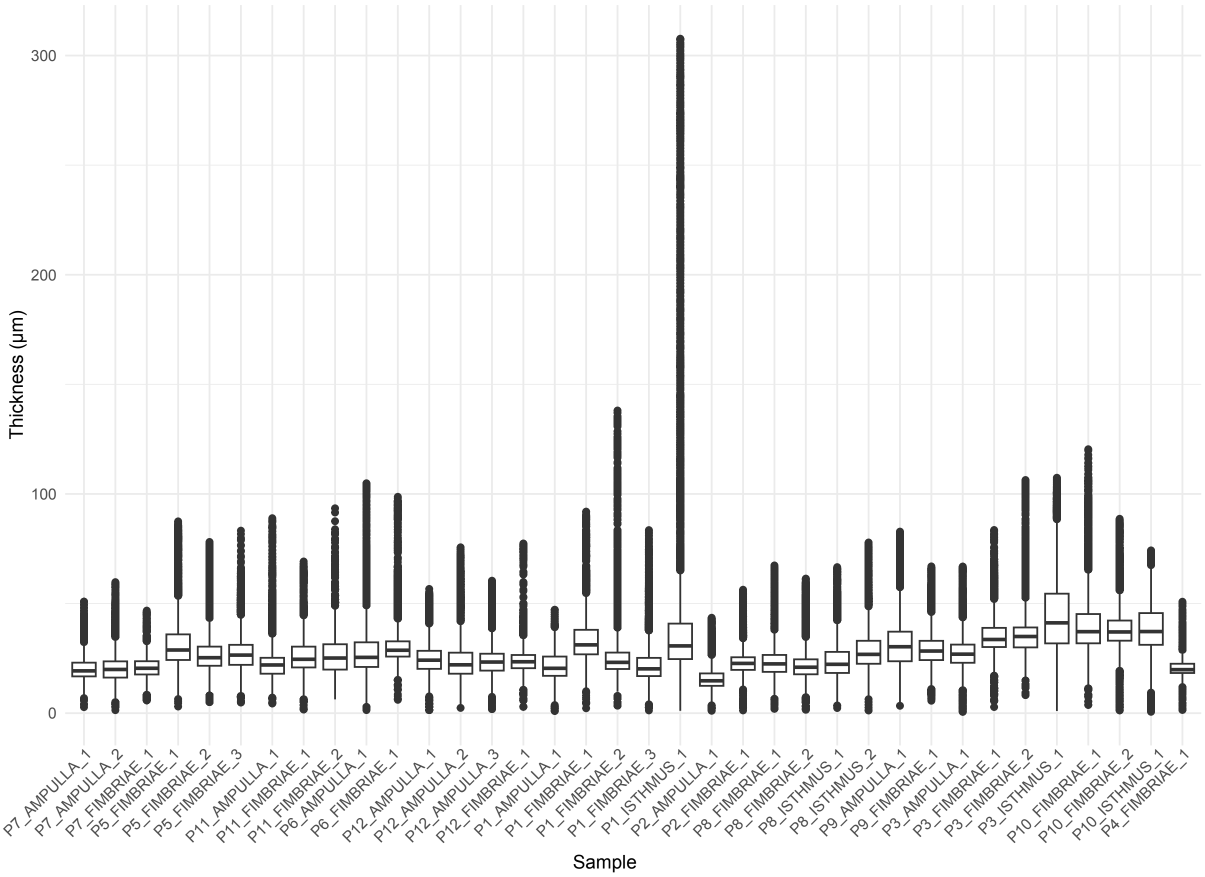


**Python measurement output.** Distribution of thickness along the measured epithelium for each sample respectively, visualizing the median thickness of each sample. Each dot represents one measuring point.
